# Rosettes in a matrix: Predicting spatial variation in density of a large felid in a forest-production mosaic

**DOI:** 10.1101/2024.06.17.599292

**Authors:** Anish Paul, Nitish Kumar, Tonmoy Mukherjee, Amir Kumar Chhetri, Aritra Kshettry

**Affiliations:** Department of Ecology and Environmental Sciences, School of Life Sciences, Pondicherry University, Puducherry, India; Post-Graduate Program in Wildlife Biology and Conservation, National Centre for Biological Sciences-TIFR, Bangalore, India; The Coexistence Project, Malbazar, India; Wildlife Conservation Society-India, Bangalore, India; Centre for Wildlife Studies, Bangalore, India; World Wide Fund for Nature (WWF) India, New Delhi, India

**Keywords:** Conservation-Compatible-Landscapes, *Panthera pardus*, human-use landscapes, tea plantations, spatially explicit capture-recapture

## Abstract

Large carnivores are keystone for ecosystems and flagships for conservation efforts but face severe threats globally. Protected Areas are vital for the conservation of these charismatic species along with a host of ecological processes. However, the extent and scope of protected areas for conservation of all threatened species is limited, especially in the global south. Considering larger landscapes that can be compatible with large carnivore conservation goals is an alternative approach to ensure their persistence. This study explores the potential of multi-use landscapes for the persistence of a globally threatened large felid, the leopard. This study investigated the spatial variability of leopard densities across a land-use gradient using spatially explicit capture-recapture framework in a tea-plantation dominated forest-production landscape mosaic. While the density of leopards in this landscape was estimated to be 7.96 ± 1.56 (SE) per 100 km^2^, significant (*p*=0.048, t=2.02, df=61) differences in estimates were observed between tea-plantations (11.53 ± 2.72 (SE) leopards per 100 km^2^) and the forested habitats (4.67 ± 2.07 (SE) per 100 km^2^). Densities between tea plantations and Protected Areas (a subset of the forested habitat) were found to be comparable (9.19 ± 4.55 (SE) per 100 km^2^). The study posits that conservation-compatible land use in landscapes shared with people can host a higher density of large felids than forested areas and that conservation planning needs to move beyond the dominant PA-centric paradigm. The study also reinforces the importance of multi-use landscapes for wildlife conservation, especially for an adaptable large felid.

## 1 Introduction

Large-bodied felids, owing to their charismatic nature, ecological role and profound cultural veneration, have always been under immense research focus (Chaudhary et al., 2019). They play vital roles in maintaining the ecosystem’s health and function as flagship species of their habitat (Ripple et al., 2014). Over the past century of the Anthropocene, many large-bodied carnivore species have experienced a drastic reduction in their global population and distribution (Ripple et al., 2014). Until recently, it was believed that large-bodied carnivores, to thrive and survive, need isolated suitable natural habitats devoid of any human interventions (Carter & Linnell, 2016; Woodroffe, 2000). Conventional conservation and wildlife management planning revolved around earmarking a piece of habitat as a protected area for wildlife to roam around without considering any possibility of human-wildlife co-occurrence in shared spaces (Woodroffe, 2000). However, protected areas account for only 12% of the world’s total land area (Chape et al., 2005) and even less in developing countries (Ghosh-Harihar et al., 2019). Even though PAs play a crucial role in maintaining population of endangered animals, they fail to conserve large-bodied species if the area is too small to meet the ecological needs of wide-ranging species (Brashares et al., 2001; Ghosh-Harihar et al., 2019; Packer et al., 2013). Moreover, increasing developmental pressures on Protected Areas, especially in the global south further threaten the paradigm of protected areas as inviolate parks (Ghosh-Harihar et al., 2019). Conversely, the PA-centric conservation paradigm automatically excludes shared landscapes that hold the potential to conserve biodiversity along with human lives and livelihoods.

Production landscapes with appropriate management regimes can host biodiversity beyond the protected area boundaries (Ghosh & Basu, 2022; Wright et al., 2012). There is increasing evidence of the survival and persistence of carnivores in human-modified multi-use landscapes across their global distribution (Alexander et al., 2016; Bateman & Fleming, 2012; Kshettry et al., 2020; Majgaonkar et al., 2019; Riley et al., 2021; Schuette et al., 2013; Valeix et al., 2012; Warrier et al., 2020; Wilmers et al., 2013). While it creates opportunities for wildlife conservation, it presents challenges in terms of spatial and temporal overlap between humans and wildlife that can lead to negative interactions between the two. Hence, carnivore conservation now calls for a landscape-level approach by considering spaces beyond protected area boundaries while proposing wildlife management plans that consider larger landscapes that are shared with people. However, this approach is still largely missing from traditional wildlife and conservation management plans. Moreover, the current ecological knowledge about large carnivores is primarily based upon the information gathered from studies conducted within protected areas, and, therefore, may be inadequate to propose carnivore management strategies outside PAs (Ghosal et al., 2013).

Globally, several large carnivore species have increased their range in the recent past and are distributed well beyond the boundaries of Protected Areas (Chapron et al., 2014; López-Bao et al., 2017). Mountain lions (*Puma concolor*) and leopards have been known to persist in densely populated peri-urban and urban areas (Riley et al., 2021; Surve et al., 2022). Even with a high density of humans, production landscapes in India have been known to provide refuge for large-bodied carnivores (Athreya et al., 2013; Kshettry et al., 2020; Warrier et al., 2020). Studies on leopards from human-use areas have revealed that in the presence of abundant prey base, vegetation cover, and compatible land-use practices, leopards can even persist in areas with high human densities (Jacobson et al., 2016; Kshettry et al., 2020). However, the relative importance of natural and shared (human-dominated) landscapes has rarely been investigated for large carnivores leading to a gap in knowledge on how population densities vary across such landscapes. Understanding such variation in densities across shared landscapes and ‘natural habitats’ for adaptable large carnivores could have widespread implications for their conservation as well as to devise appropriate conflict mitigation measures. The leopard is an ideal model species to study the effects of land use on its density owing to its widespread distribution and adaptability (Jacobson et al., 2016). This study attempts to understand the variation in leopard density across a land-use gradient to understand the relative importance of natural habitats (with varying protection regimes) and habitats shared with people. The paper also discusses the need to balance human safety and carnivore conservation in shared landscapes for equitable and inclusive conservation models.

In India, leopards mostly share their space with the other charismatic large carnivore, the Bengal tiger (*Panthera tigris tigris*), in most protected areas (Chaudhary et al., 2019). Hence, most of the information available about the ecology of leopards is a bycatch of studies focused on tigers, and they mainly deal with the estimation of abundance (Jhala et al., 2008; Harihar et al., 2011; Kalle et al., 2011). Very few studies have been focused on the species ecology (but see Borah et al., 2013; Mandal et al., 2017; Noor et al., 2020; Surve et al., 2022; Thapa et al., 2014), and fewer attempted to study their ecology in human-modified multi-use landscapes (but see Athreya et al., 2016; Gubbi et al., 2020; Kshettry et al., 2018, 2020; Majgaonkar et al., 2019; Naha et al., 2021; Sidhu et al., 2017). However, no study has attempted to understand the variation in leopard densities across land-use gradients from the same landscape thereby leaving a lacuna which has been addressed by the current study which was carried out in a forest-production landscape mosaic.

The Duars region of northern West Bengal represents a heterogeneous matrix of protected areas, reserve forests, tea estates, agricultural fields, human settlements, and waterbodies. The region has been proven to be extensively used by leopards to reside, reproduce, and hunt (Kshettry et al., 2017, 2020; Naha et al., 2020). Thus, the landscape was deemed appropriate to investigate the habitat factors influencing the species density in this shared landscape. The study also provides a framework for estimating carnivore population density in shared landscapes through incorporating the knowledge and experiences of local communities and stakeholders into camera-trapping methodology. The study investigates the relative importance of natural and shared spaces for conserving a threatened and charismatic felid by through estimating their densities across a land-use gradient.

## 2 Article types

Original Research

## 3 Materials and Methods

### 3.1 Study Area

The study was conducted in an area of ∼400 km^2^ situated in Duars region of northern West Bengal (Figure 1). The region constitutes a portion of the East-Himalayan Biodiversity Hotspot and is historically represented by naturally occurring Sal (*Shorea robusta*) and Simul (*Bombax ceiba*) dominated forests (Champion & Seth, 1968; Myers et al., 2000). The region is represented by two major forest types: Northern Tropical Semi-evergreen forests and Tropical Moist Deciduous forests, along with some remnant patches of Eastern Alluvial Grasslands (Bhattacharjee & Parthasarathy, 2013; Roy, 2014). Apart from naturally occurring vegetation, artificial monoculture plantations of several economically important species like sal (*Shorea robusta*), teak (*Tectona grandis*), and rubber (*Ficus elastica*) are present in a patchy distribution within the entire landscape (Roy, 2009). During the late 1800s, vast stretches of these forests were converted to tea plantations by the British Colonials (Kshettry et al., 2018). Presently, the remaining forest patches stand isolated, interspersed with numerous tea estates, agricultural fields, and human habitations (Kshettry et al., 2020). Currently, a significant portion of the naturally occurring forests is represented by two protected areas, namely Chapramari Wildlife Sanctuary (9.5 km^2^) and Gorumara National Park (80 km^2^), and the rest falls under the Reserve Forests of Jalpaiguri Territorial Division. The Protected Areas and Reserve Forests in the study area have differing protection and human activity regimes and hence, were included as two separate classes of ‘natural habitats’ for the leopard density estimation. The Protected Areas have lower human activity and higher habitat management and wildlife conservation efforts compared to the reserve forests.

**Figure 1:**
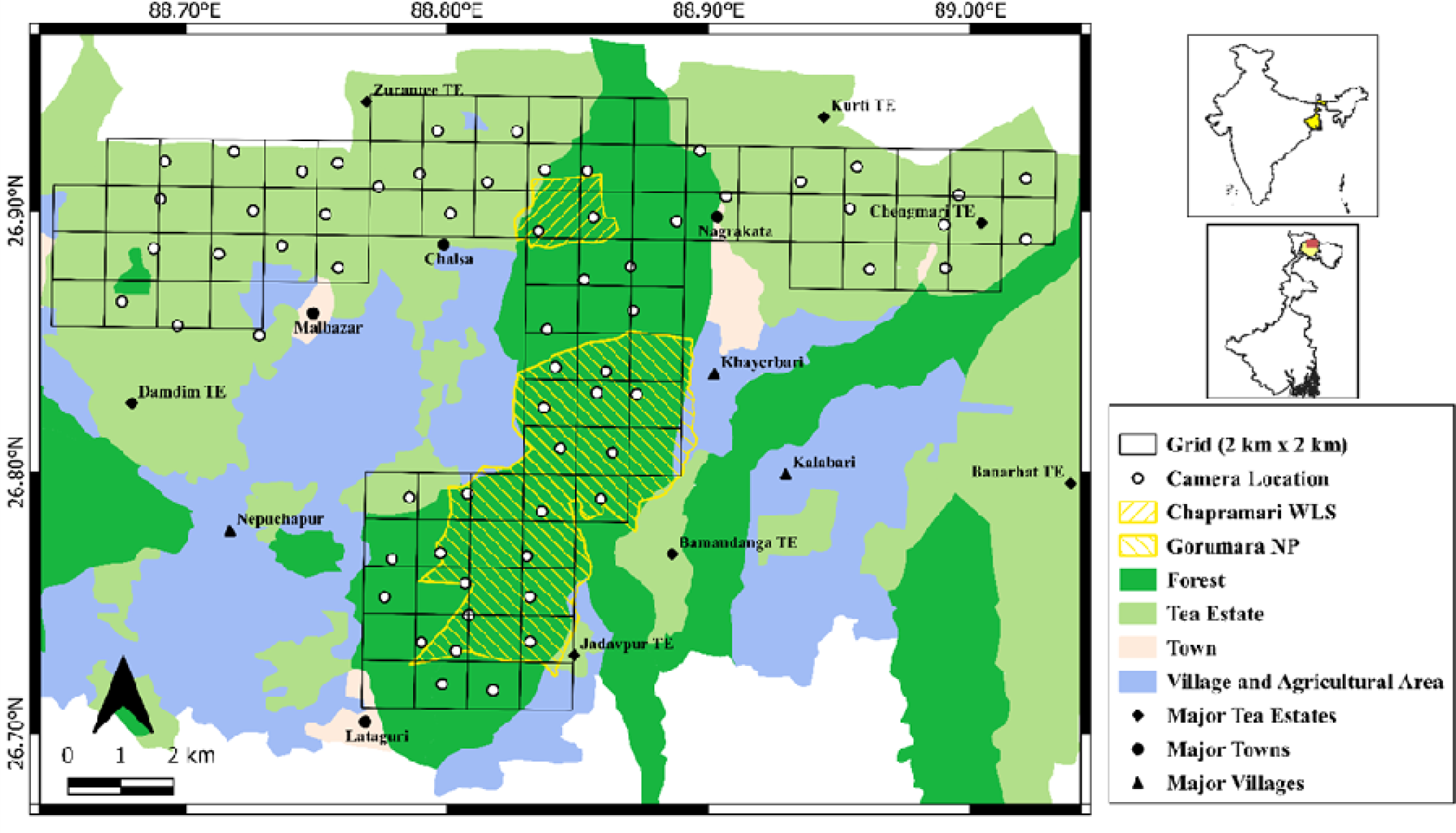
A map of the study area depicting the placement of camera traps within the forest-tea estate mosaic landscape.

As per the 2011 census report, the region beholds a population density of 701 persons per km^2^ (Kshettry et al., 2017). The daily per-capita income of people is often less than 1 USD, and agriculture and livestock rearing constitute the most predominant source of livelihood apart from tea production (Kshettry et al., 2017). Leopard is the apex predator of the landscape and readily uses both forests and tea estates as its habitat (Kshettry et al., 2017; Naha et al., 2021). Other major wild species from the area include Asian elephant (*Elephas maximus*), gaur (*Bos gauras*), one-horned rhinoceros (*Rhinoceros unicornis*), sambar (*Rusa unicolor*), chital (*Axis axis*), muntjac (*Muntiacus muntjac*), rhesus macaque (*Macaca mulata*), wild boar (*Sus scrofa*), jungle cat (*Felis chaus*), leopard cat (*Prionailurus bengalensis*) and Indian peafowl (*Pavo cristatus*) (Bhattacharjee & Parthasarathy, 2013). These species constitute the wild prey base for the leopards (Kshettry et al., 2018).

### 3.2 Data Collection

Owing to their cryptic nature and low population density, it is tough to deduce an unbiased population estimate for large carnivores (Boulanger et al., 2004; Karanth, 1995). For cryptic carnivores, camera traps help to infer an almost accurate density estimate (Balme et al., 2009). Carnivore population estimation using the photographic capture-recapture technique was first implemented in India for tigers in Nagarhole National Park (Karanth, 1995). Since then, with the slightest of modifications, this method has been widely used to estimate densities of animals bearing distinct natural markings like tiger (*Panthera tigris)* (Karanth et al., 2006; Rather et al., 2021), lion (*Panthera leo*) (Elliot & Gopalaswamy, 2017; Gogoi et al., 2020), snow leopard (*Panthera uncia*) (Alexander et al., 2015; Sharma et al., 2021), jaguar (*Panthera onca*) (Maffei et al., 2004; Silver et al., 2004), leopards (*Panthera pardus*) (Athreya et al., 2013; Harihar et al., 2009; Surve et al., 2022), ocelots (*Leopardus pardalis*) (Maffei & Noss, 2008; Trolle & Kéry, 2003), hyena (*Hyaena sp.*) (Singh et al., 2014) as well as of species which lack distinct natural marks, such as rusty-spotted cat (*Prionailurus viverrinus*) (Chatterjee et al., 2020) and dhole (*Cuon alpinus*) (Punjabi et al., 2022).

The entire study area was divided into grid cells of 2 km x 2 km. For forest patches, locations were identified based on preliminary sign surveys (Karanth & Nichols, 1998), prior knowledge of the site and information from the forest officials. Preliminary sign surveys were also conducted to identify all possible trails frequently used by leopards for the tea estates. Due to heavy anthropogenic traffic, it becomes hard to detect animal signs in tea estate trails. Hence, the tea estate authorities, workers and residents were consulted to identify the areas of frequent leopard sightings. All levels of the tea estate management were made aware of the exercise to obtain full cooperation. Some residents and the guards of the estates were involved to ensure the safety and security of the cameras. The dry season in the study area overlapped with the pruning season of the tea bushes leading to the loss of vegetation cover, which brings in fewer leopard activities in the pruned portions (Kshettry et al., 2017; Naha et al., 2021). Hence, sections earmarked for pruning were avoided while selecting camera locations. Each site was provided with two camera traps deployed opposite each other to obtain photographs of both the flank of the animal (Karanth & Nichols, 1998). Cameras deployed within the forests were kept operational for 24 hours daily and checked every 7-10 days. Camera traps operating within tea estates were attended to daily and were removed at 06:00 hrs from the location and reinstalled at 17:00 hrs at the same place to avoid damage and theft (Athreya et al., 2013; Surve et al., 2022).

All the photographs were tagged and classified manually using Digikam 7.0.0 software. To maintain homogeneity, only the photographs captured in forests within the time window of 06:00-17:00 were used for analysis. Photographs with any distortions or lack of clarity were discarded from any further use (Karanth & Nichols, 1998). All the photo-captured leopards were identified manually, and a capture-history file was made. Due to technical issues, both flanks of a few individuals could not be obtained. Hence, only right-flanked images were used to primarily identify individuals as it was higher in the number unless the left flank matched with a previously photographed leopard whose both flanks were profiled (Athreya et al., 2013; Harihar et al., 2009).

### 3.3 Analysis

Each trap night was treated as an independent sampling occasion (Otis et al., 1978). Data from all four blocks were pooled together to construct individual capture histories. The cumulative number of leopard individuals captured was plotted against the camera trap occasions to check for sampling adequacy. Successive trap movement of leopard individuals was extracted from the raw photo-captures using capture history and was plotted to see the extent of movement by leopards between traps. The analysis was done using the package “secr” (Efford 2023) v4.5.10 in Program R (R Core Team, 2018). The “suggest.buffer” function was used to create a habitat mask around the trap sites over which the density would be estimated (Sharma et al., 2021). This function uses the movement patterns of individual leopards from the capture history to predict a buffer width which ensures that any animal having activity centres beyond that distance from the outermost trap will have a negligible probability of getting photo-captured in any of the detectors. All the non-habitat areas, like towns and villages, were removed from the mask layer. Agricultural lands, although known to constitute habitats for large carnivores in parts of the country (Warrier et al., 2020), were also removed from the final mask layer as during the study period, there were no standing crops in these areas to provide cover to the leopards. Furthermore, previous study in the landscape found low habitat-use probability of leopards in agriculture fields in the region (Kshettry et al., 2020).

Null models (D∼1, g0∼1, sigma∼1) were fitted with three different detection functions, i.e., Half Normal (HN), Negative Exponential (EX) and Hazard Rate (HR). Among the null models, four models were tested for the study, i.e., null model (D∼1, g0∼1, sigma∼1), global learned response model (D∼1, g0∼b, sigma∼1), site-specific learned response model (D∼1, g0∼bk, sigma∼1) and 2-part individual heterogeneity model (D∼1, g0∼h2, sigma∼1).

### 3.4 Covariate Development

The leopard population density in the study area was modelled as a function of ecological and anthropogenic covariates. Covariates were chosen based on the available literature and were hypothesized to have different expected relations with leopard density (Table 1). A habitat mask layer of ∼6 km was prepared using the “secr” package (Efford 2023) v4.1.2 in R (R Core Team, 2018). All the points of the mask layer were given their land use-land cover identity, and the non-habitat points were removed to prepare the final mask layer (Supplementary Figure S2).

**Table 1.** Habitat Covariate Development and Their Expected Relations with Leopard Density and Factor It Is Hypothesized to Influence.

| Covariates | Source | Expected Relation with Density | Hypothesized Influenced Parameter | Reference |
| --- | --- | --- | --- | --- |
| Habitat Type (HabType) | Remotely Sensed | +/- | Density (D) | Alexander et al., 2016; Havmøller et al., 2019; Rather et al., 2021 |
| Distance from Waterbody (DisWB) | Remotely Sensed | - | Density (D) | Naha et al., 2021; Rather et al., 2021 |
| Distance to nearest Human Settlement (DisHS) | Remotely Sensed | + | Density (D) | Alexander et al., 2016; Havmøller et al., 2019; Naha et al., 2021 |

We tested the influence of five covariates on the detection probability (g0). Detection probability (g0) was hypothesized to be a function of Euclidian distance to nearest waterbody, nearest human settlement and nearest road from each camera trap location. Proportions of the total operational trap night of each camera trap location on which humans and prey species were photo-captured were also added as a covariate of detection function (g0). The influence of three covariates on leopard density was tested hypothesizing their different relationships with density (D) (Table 1). Euclidian distance of the nearest waterbody and nearest human settlement from each point of the mask layer. QGIS v3.16.15 was used to estimate all the distances. We standardized all covariate values before using them in the model.

Habitat type was added as a covariate to estimate density separately for entire forested habitat (Protected Area + Reserve Forests) and tea estates (Supplementary Figure S2). Furthermore, only the Protected Areas were included as another habitat type which was a subset of the forested habitat, and we compared the density between the tea plantations and protected forests. Akaike Information Criterion corrected for small sample size (AICc) was calculated for all the competing models, and the most suitable model in each case was selected based on the difference between the corresponding AICc values (ΔAICc) (Burnham & Anderson, 2001).

## 4 Results

### 4.1 Leopard Photo Capture

The sampling area was divided into four blocks, and the exercise was carried out from December 2021 to March 2022 in four trapping sessions of 20 days each. A total of 63 trap locations were used to sample the four blocks. Block 1 had 14 locations, Block 2 had 15 locations, Block 3 had 23 locations, and Block 4 had 11 locations. The average spacing between all the trapping sites was 1.8 km which ensured that all leopard individuals had an equal probability of getting photo-captured without leaving major holes in the trap array (Athreya et al., 2013; Balme et al., 2009; Odden & Wegge, 2005). A total of 71 photographic detections of 32 individuals were obtained during the period of 1205 trap nights. Camera traps at two locations became non-functional due to theft. Some of the cameras were damaged by elephants, while some appeared non-functional for a few nights due to technical errors. Out of 63 camera locations, 35 had leopard detections.

The number of unique individuals detected did not reach an asymptote during the study period (Supplementary Figure S3). However, studies have shown that even after a prolonged sampling effort, it is likely impossible to capture all the individuals present in a landscape (Sharma et al., 2010). Trap-revealed leopard movement based on photo captures showed that almost 77% of all movements occurred within the range of 0-3000 meters (Supplementary Figure S4). Only a negligible proportion (7.7%) of all movements occurred beyond 6000 meters. The “suggest.buffer” function in “secr” produced a buffer of 6075 meters around each camera trap. Altogether, it created a mask layer with an area of 1036.31 km^2^. All the non-habitat areas removed from the suggested mask layer finally yielded an effective sampling area of 759.64 km^2^ (Supplementary Figure S2). The effective sampling area comprised of 416.23 km^2^ of tea-plantations and 343.41 km^2^ of forested area (comprising of Reserve forests and Protected Areas. The area under Protected Area habitat (a subset of the forested area) was 82.4 km^2^.

### 4.2 Model Fitting and Density Estimation

Models were fitted under the assumption of a null model using three different detection functions, i.e., HN, EX and HR, to estimate the density. The HN model had the lower AICc score and was used for the subsequent model-building process (Supplementary Table S2, S3, Supplementary Figure S5).

The estimation of leopard density was carried out through two broad model categories. The first category includes the null model and four parameters hypothesized to influence the detection function (g0): global learned response model (D∼1, g0∼b, sigma∼1), site-specific learned response model (D∼1, g0∼bk, sigma∼1), 2-part individual heterogeneity model (D∼1, g0∼h2, sigma∼1) and time factor model (D∼1, g0∼t, sigma∼1). The estimated density of leopards using the Null model was found to be 7.96 ± 1.56 per 100 km^2^ (Mean ± SE) (Table 2). The heterogeneity model (h2) performed as the second-ranked following the null model (Table 3).

**Table 2.** Density Estimates (No./100 km2) of Leopard using null model (g0∼1), global behaviour response model (g0∼b), site-specific behaviour response model (g0∼bk), heterogeneity model (g0∼h2) and time factor model (g0∼t).

| Model | Parameters | link | estimate | SE.estimate | lcl | ucl |
| --- | --- | --- | --- | --- | --- | --- |
| null | D | log | **7.96** | **1.56** | 5.43 | 11.67 |
|  | g0 | logit | 0.038 | 0.008 | 0.025 | 0.057 |
|  | sigma | log | 1851.691 | 197.622 | 1503.076 | 2281.161 |
| b | D | log | **7.08** | **1.44** | 4.77 | 10.50 |
|  | g0 | logit | 0.053 | 0.016 | 0.029 | 0.096 |
|  | sigma | log | 1848.116 | 199.283 | 1469.955 | 2281.653 |
| bk | D | log | **8.19** | **1.66** | 5.51 | 12.16 |
|  | g0 | logit | 0.03 | 0.009 | 0.017 | 0.053 |
|  | sigma | log | 1977.193 | 246.193 | 1550.484 | 2521.335 |
| h2 | D | log | **9.19** | **2.38** | 5.57 | 15.16 |
|  | g0 (h2 = 1) | logit | 0.022 | 0.013 | 0.006 | 0.071 |
|  | g0 (h2 = 2) |  | 0.065 | 0.028 | 0.027 | 0.148 |
|  | sigma | log | 1867.162 | 202.441 | 1510.645 | 2307.819 |

Note: Three model parameters estimated are D = density per 100 km<sup>2</sup>, g0 = probability detection and sigma = distance over which detection probability gradually decreases

**Table 3.** Summary of AIC values for Density Estimates using null model (g0∼1), global behaviour response model (g0∼b), site-specific behaviour response model (g0∼bk), heterogeneity model (g0∼h2) and time factor model (g0∼t).

| model | detectfn | npar | logLik | AIC | AICc | dAICc | AICcwt |
| --- | --- | --- | --- | --- | --- | --- | --- |
| D~1 g0~1 sigma~1 | halfnormal | 3 | -161.1481 | 328.296 | 329.153 | 0.000 | 0.7721 |
| D~1 g0~h2 sigma~1<br>pmix~h2 | halfnormal | 5 | -159.6429 | 329.286 | 331.593 | 2.440 | 0.2279 |
| D~1 g0~bk sigma~1 | halfnormal | 4 | -359.9167 | 727.833 | 729.315 | 400.162 | 0.0000 |
| D~1 g0~b sigma~1 | halfnormal | 4 | -359.8814 | 727.763 | 729.244 | 400.091 | 0.0000 |

We fitted five different models for detection function (g0) while keeping the density constant. None of the parameters influenced the detection function significantly (Supplementary Table S4). Hence the null model of detection function (g0) was retained following the principle of parsimony. Once the detection model was fixed, the density model was built using this fixed detection model.

In the second model category, the density (D) was considered to be influenced by additive effects of three habitat covariates-Habitat Type, Distance to Waterbody, and Distance to Human Settlements. AICc scores comparison suggested the top model for density to be a function of both Habitat Type and Distance to Human Settlements, followed by the one with Habitat Type alone (Table 4). The model with Habitat Type as a covariate of density revealed that leopard density was highest in tea estates with an estimate of 11.53 ± 2.72 per 100 km^2^ (Mean ± SE). Estimated density for the forested area was 4.67 ± 2.07 per 100 km^2^ (Mean ± SE) (Supplementary Table S5, Figure 2). Density for the Protected Areas (a subset of the forested habitat) was estimated to be 9.19 ± 4.55 per 100 km^2^ (Mean ± SE).

**Figure. 2.**
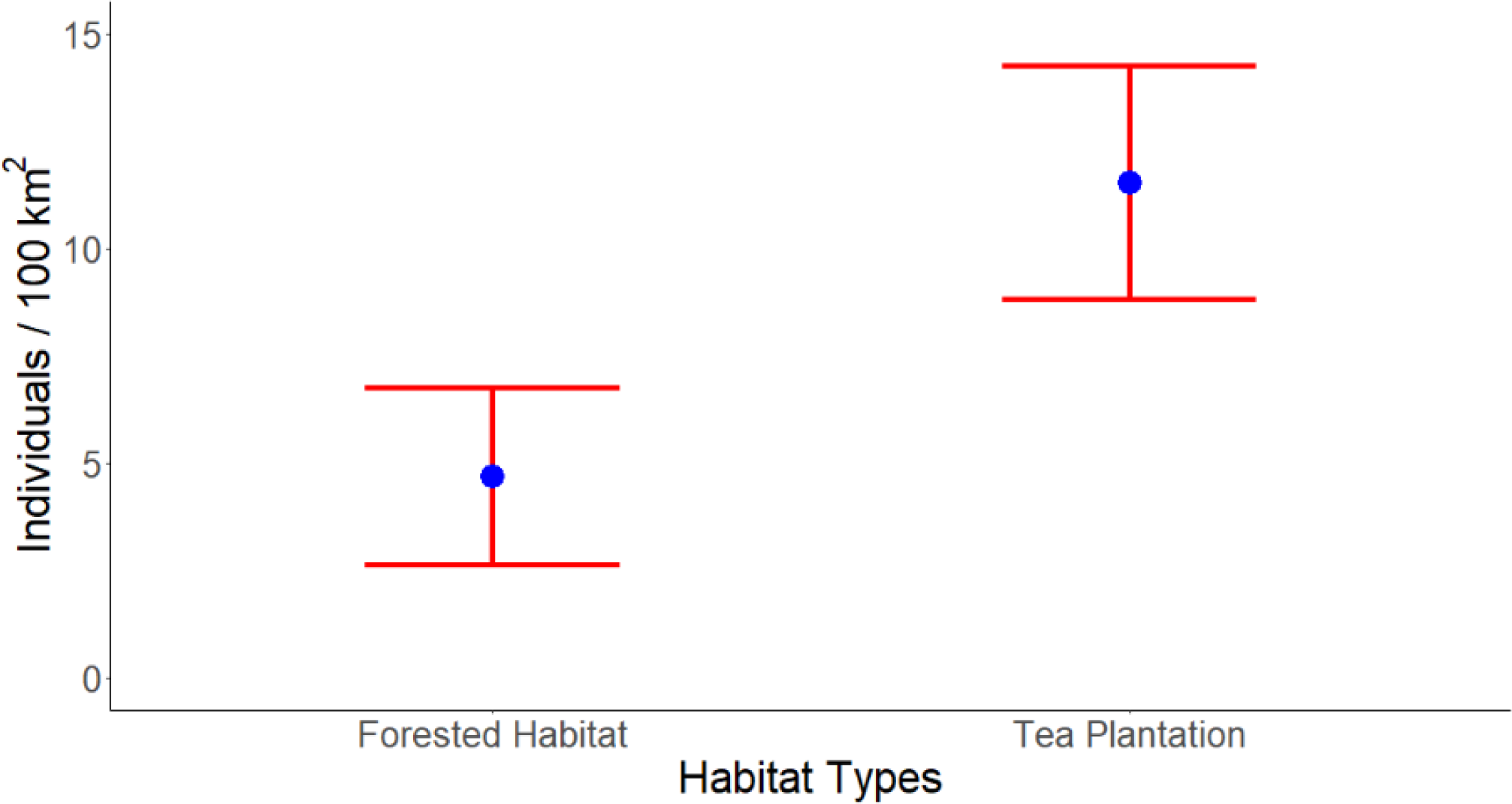
Density estimates / 100 km^2^ (Mean+SE) of Indian leopards in different habitat and management types. Red whiskers represent the standard error around the mean.

**Figure. 3.**
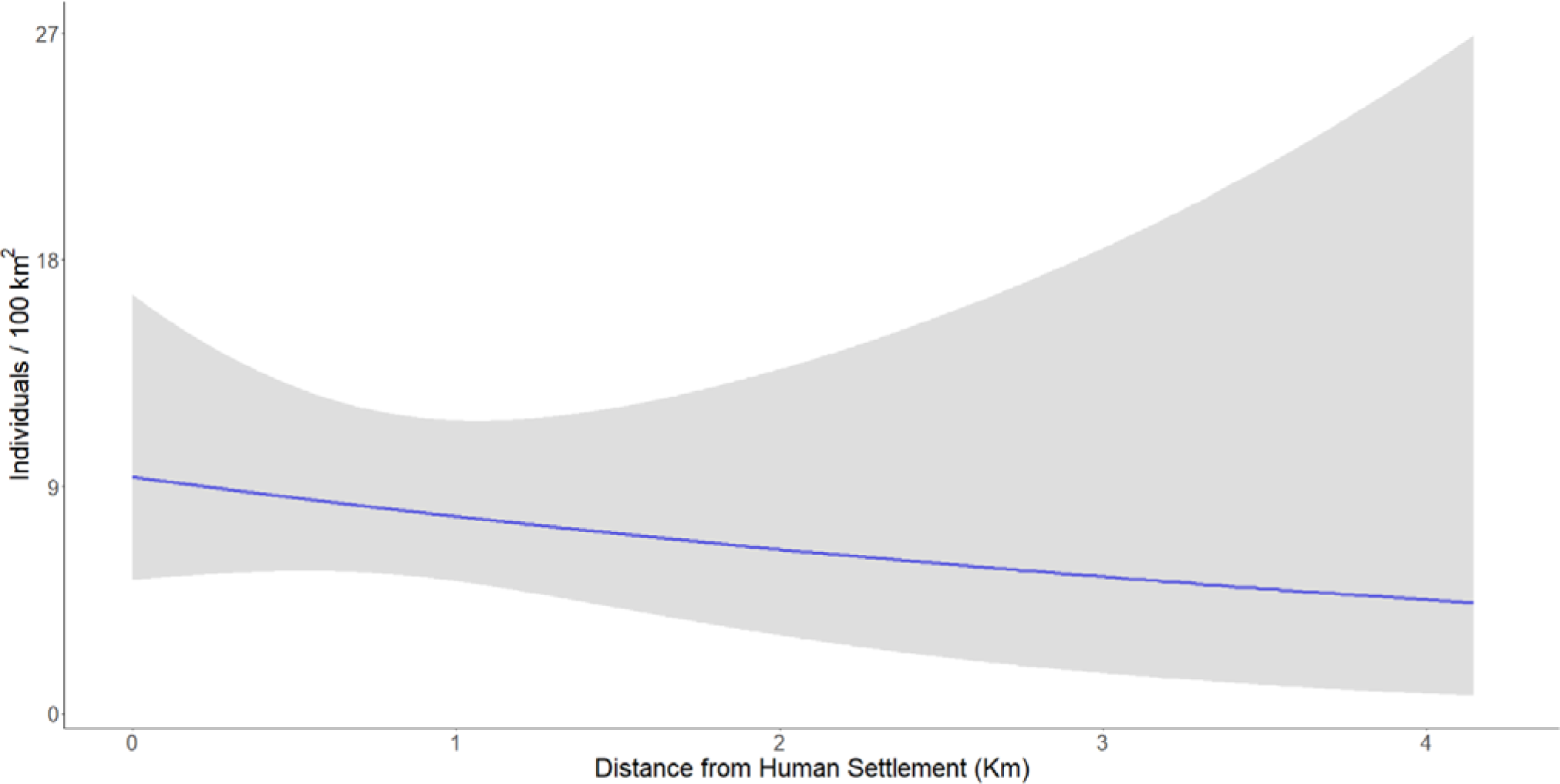
leopard density as a function of distance to nearest human settlements. Gray shaded area represents the 95% confidence intervals.

**Table 4.** AICc based model selection for models with leopard density as a function of different covariates.

|  | Model | detectfn | npar | logLik | AIC | AICc | dAICc | AICcwt |
| --- | --- | --- | --- | --- | --- | --- | --- | --- |
| Habitat Type +<br>Distance to Human<br>Settlement | D~HabType + DisHS<br>g0~1 sigma~1 | halfnormal | 5 | -157.1902 | 324.380 | 326.688 | 0.000 | 0.4107 |
| Habitat Type | D~HabType g0~1<br>sigma~1 | halfnormal | 4 | -159.5617 | 327.123 | 328.605 | 1.917 | 0.1575 |
| Distance to Waterbody | D~DisWB g0~1 sigma~1 | halfnormal | 4 | -159.7859 | 327.572 | 329.053 | 2.365 | 0.1259 |
| Null | D~1 g0~1 sigma~1 | halfnormal | 3 | -161.1481 | 328.296 | 329.153 | 2.465 | 0.1197 |
| Global | D~HabType + DisWB +<br>DisHS g0~1 sigma~1 | halfnormal | 6 | -157.1211 | 326.242 | 329.602 | 2.914 | 0.0957 |
| Habitat Type +<br>Distance to Waterbody | D~HabType + DisWB<br>g0~1 sigma~1 | halfnormal | 5 | -159.3184 | 328.637 | 330.945 | 4.257 | 0.0489 |
| Distance to Human Settlement | D~DisHS g0~1 sigma~1 | halfnormal | 4 | -160.8896 | 329.779 | 331.261 | 4.573 | 0.0417 |

Note: HabType = Habitat type, DisHS = distance to nearest human settlement, DisWB = distance to nearest waterbody. Three model parameters are D = density per 100 km<sup>2</sup>, g0 = probability detection and sigma = distance over which detection probability gradually decreases

**Table 5.** Estimated parameters from the top-ranking model of density estimation.

|  | **beta** | **SE.beta** | **lcl** | **ucl** |
| --- | --- | --- | --- | --- |
| D | -10.152388 | 1.2157899 | -12.5352925 | -7.769484 |
| D.HabTypeTE | 3.891780 | 1.3051192 | 1.3337935 | 6.449767 |
| D.DisHS | 1.133706 | 0.3535931 | 0.4406767 | 1.826736 |
| g0 | -3.288037 | 0.2202218 | -3.7196641 | -2.856411 |
| sigma | 7.550262 | 0.1080556 | 7.3384770 | 7.762047 |

Note: HabType = Habitat type (Forest and Tea Estate), DisHS = distance to nearest human settlement. Three model parameters are D = density, g0 = probability detection and sigma = distance over which detection probability gradually decreases

## 5 Discussion

We estimated leopard densities from a tea-plantation dominated landscape of north-eastern India and investigated the reasons for the spatial variability in density. The study also underscored including local knowledge and collaboration for carrying out such large-scale exercises in human-dominated landscapes. Our density models indicated that variation in the leopard density is largely predicted by the habitat types/land-use types present in the landscape. However, the top model included distance to human settlements, confirming the presence of patterns within each habitat type. Distance to human settlements was found to be negatively correlated with leopard density. This can be an outcome of the high prey availability near the human settlements in the form of livestock. Previous studies from the landscape have shown that domestic prey density is six times higher than wild prey availability, and 60% of the prey biomass of the leopard diet comprises domestic prey species (Kshettry et al., 2018).

Habitat-specific density estimate showed leopard density in the tea estate to be 11.53 ± 2.72 leopards per 100 km^2^ (Supplementary Table S5). The only other leopard density estimation study from a human-dominated multi-use landscape reported a density of 4.8 ± 1.2 individuals per 100 km^2^ (Athreya et al., 2013) which is much less than the current estimate. The current estimate of leopard density in tea estates of the study area is even greater than the same from many of the protected areas of the country (Figure 4), such as Bandhavgarh Tiger Reserve (Rather et al., 2021), Dachigam National Park (Noor et al., 2020), Manas National Park (Borah et al., 2013), Pakke Tiger Reserve (Selvan, 2013), Rajaji Tiger Reserve (Harihar et al., 2011), Sariska Tiger Reserve (Mondal et al., 2012) and Tungareshwar Wildlife Sanctuary (Surve et al., 2022). Only three other protected areas in the country, Mudumalai Tiger Reserve (Kalle et al., 2011), Achanakmar National Park (Mandal et al., 2017) and Sanjay Gandhi National Park (Surve et al., 2022) have been known to have a comparable or greater density of leopards than this estimate. However, this comparison is relevant only to underscore the conservation potential of a production landscape mosaic for leopards and exploring the reasons for this variation in densities is beyond the scope of this study. However, we can speculate that the variation in densities discussed in this and subsequent section can be a function of the variety of habitats in these areas, prey availability, and the presence of sympatric carnivores. Even within the study area, spatial variation in leopard density was observed across land-use types.

**Figure. 4.**
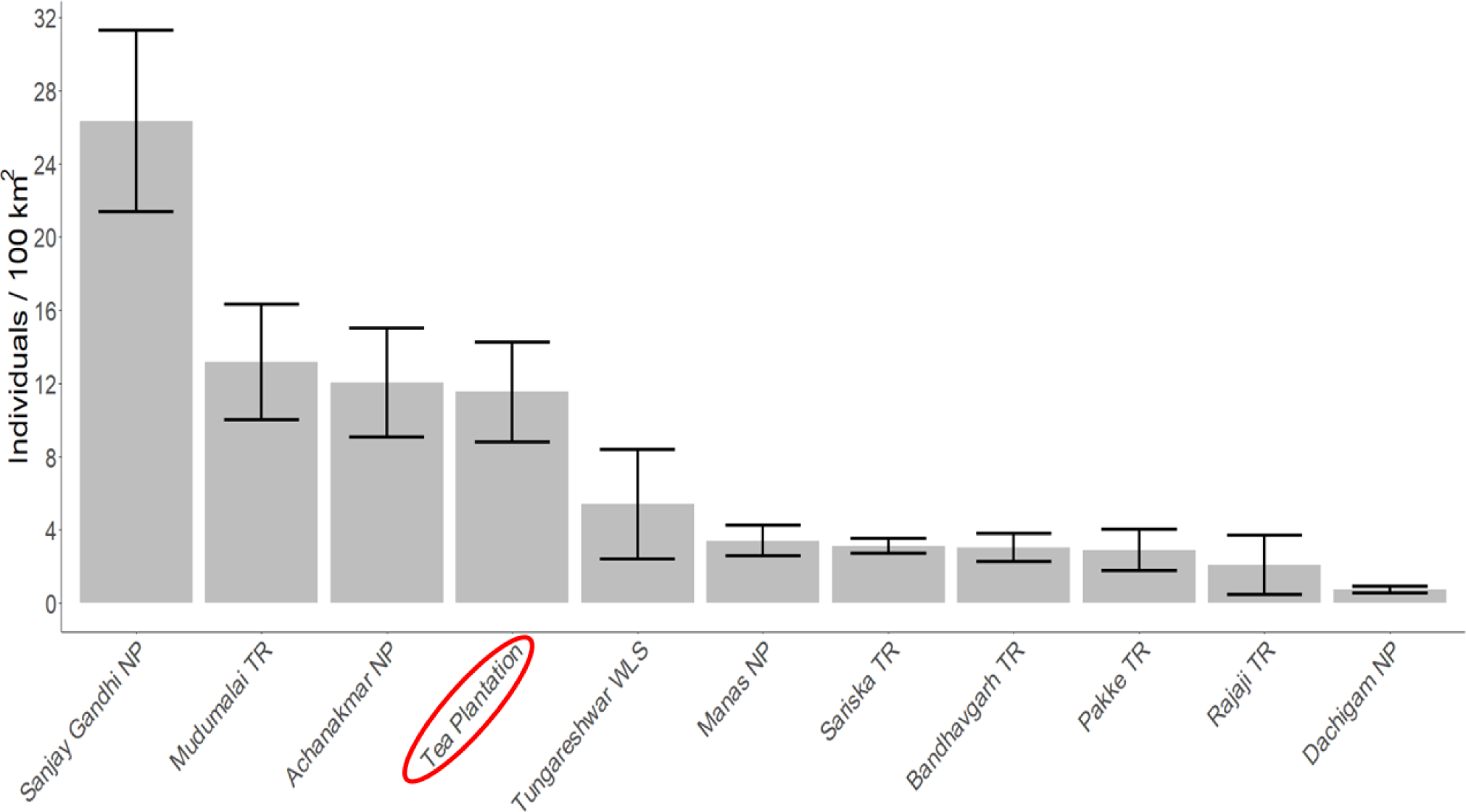
Comparison of leopard density from camera-trap studies conducted in different protected areas across India using spatially-explicit capture-recapture framework. Bars represent the density estimates. Black whiskers represent the standard error around the mean. The red ellipse indicates the estimate from the tea plantation habitat in the current study area.

Density of leopards within the forested areas (reserve forests and Protected Areas) was significantly lower that the tea plantations. This could be speculated to be the result of very low prey densities in the reserve forest areas. Studies have reported that reserve forests of the region experience a greater extent of deforestation, greater anthropogenic pressure and scarcity of prey animals, which can make them incapable of supporting a substantial population of the large carnivore or herbivores (Nagendra et al., 2009). Despite that, the density is still greater than the estimates reported from Dachigam National Park (Noor et al., 2020), Manas National Park (Borah et al., 2013), Pakke Tiger Reserve (Selvan, 2013), Rajaji Tiger Reserve (Harihar et al., 2011), Sariska Tiger Reserve (Mondal et al., 2012) and Bandhavgarh Tiger Reserve (Rather et al., 2021).

The higher density of leopards in this landscape can be attributed to the absence of tigers. Studies suggest that leopards tend to occur at greater density in the absence of tigers, which often diminishes once the tiger is reintroduced (Harihar et al., 2011; Mondal et al., 2012). However, one of the country’s highest estimates of leopard density has been reported from a landscape where tigers and leopards co-occur (Kalle et al., 2011). Similarly, the lowest leopard density estimate has been reported from a region where tiger does not occur, though another sympatric species, the snow leopard (*Panthera uncia*), is present in the landscape (Noor et al., 2020).

Another reason for such a high density of leopards can be the availability of prey in the study site. Large-bodied carnivores can survive in areas closer to high anthropogenic disturbance if the prey availability is sufficient (Alexander et al., 2016). In human-dominated areas, if the availability of wild prey species is limiting, leopards are known to prey on livestock and free-ranging dogs, which is one of the reasons that makes them capable of inhabiting human-use areas (Athreya et al., 2016; Kshettry et al., 2018; Shehzad et al., 2015). Previous studies from the same landscape showed that livestock availability in the tea plantations is up to seven times higher than wild prey availability inside the forests (Kshettry et al., 2018). Estimating the relative availability of prey using their detection in the camera traps was beyond the scope of the current study due to the inability to keep the cameras operational in the tea estates during the daytime when livestock are most active. Moreover, this high density of leopards is also a function of local acceptance towards the species, a strong legislature that accords them high protection irrespective of land use and the ecological history of the landscape (Kshettry et al., 2020). Studies conducted elsewhere in India have also confirmed leopards (and other big cats) to be part of human imagination and culture and hence explain how people can share space with this charismatic felid (Dhee et al., 2019; Nair et al., 2021). However, the presence of a large felid in these shared landscapes is not without significant challenges to local lives and livelihoods.

The forest-production landscape mosaic of the Duars region in West Bengal reports one of the highest frequencies of human-leopard negative interaction globally (Kshettry et al., 2020). The tea estates of this region, which leopards use as their habitats, have been reporting incidents of human injuries due to leopards for a long time, and the numbers are ever-increasing (Bhattacharjee & Parthasarathy, 2013; Kshettry et al., 2017). During 1990-1997, there were 121 reported incidents of leopard attacks on humans, whereas, between 2001-2008, 243 incidents of humans being injured by leopards were reported from the region (Bhattacharjee & Parthasarathy, 2013). During 2009-2018, an average of 56 cases of human injury by leopard per year was reported from the area (Kshettry et al., 2020), and the spatiotemporal pattern of the attacks showed that most of such attacks took place in the tea estates during the dry season, i.e., December-April (Kshettry et al., 2017). During this time the vegetation cover in tea estates is partially lost due to the pruning of the tea bushes, and tea workers carry out intensive work within the small tea-bearing sections where the leopards also rest during the day (Kshettry et al., 2017). Livestock loss has also been known to be high in the landscape thereby incurring additional costs to local communities (Naha et al., 2020). However, despite heavy losses the communities have strong cultural and religious reverence towards local wildlife, the people of northern West Bengal still do not practice persecution of leopards as retaliatory killings of leopards are infrequent (Naha et al., 2021). The elephant too, despite causing widespread economic losses and high human casualties are worshipped by many communities in the landscape who even do not claim ex-gratia payments for their losses (Kshettry et al., 2021).

The high leopard densities in the tea-plantation habitat indicate the tremendous conservation potential of the species in such habitats and present conservation opportunities as well as challenges. In the case of our study area, the effective sampling area under tea was approximately 416.23 km^2,^ and with an average leopard density of 13 per 100 km^2^, the area could host more than 50 leopards. The region has approximately 1000 km^2^ under tea and the combined tea plantation area in West Bengal and Assam is more than 4000 km^2^, which dwarfs the area under formal protection in the region. There are also reports of leopard presence in tea estates of southern India (Navya et al., 2014; Sidhu et al., 2017) and Sri Lanka (Kittle et al., 2012). Similar studies in these regions will also help to assess the potential of tea plantations as conservation landscapes for highly adaptable species like leopards. Hence, the conservation potential of such landscapes is immense and, yet, is outside the purview of traditional conservation and management plans (Kshettry et al., 2020). Furthermore, large felids such as jaguars, lions, puma and snow leopards are also known to persist in shared landscapes and investigating the variability in densities in shared landscapes could throw up similar patterns as well thereby underscoring the conservation potential for these species. Our study attempts to take a small step towards highlighting that species conservation plans even for large carnivores can consider shared landscapes but needs to be planned in a socially just and equitable manner. Balancing conservation goals with local safety and food security can enable space sharing in such Conservation Compatible Landscapes (CCL) and needs to be the primary focus for all relevant stakeholders (Kshettry et al., 2020). The first step towards identifying such landscapes would need the estimation of the population of the species of interest and the social carrying capacity for the species.

## 6 Conclusion

The present study tries to evaluate the potential of a highly populated production landscape in supporting an apex carnivore population. The study is only the second attempt in India after Athreya et al., (2013) to study the density of leopards in a human-dominated landscape and the first ever to study leopard density in a tea-plantation landscape. The study reveals that human-modified landscapes can host significantly higher population densities of certain large carnivores than forests, even the protected areas. However, this pattern does not disregard the importance of protected areas for a host of other biodiversity and ecosystem processes, rather highlights the conservation potential of shared landscapes for global carnivore conservation. Nevertheless, the persistence of a large carnivore in a shared landscape with a very high density of humans can lead to negative outcomes for people and wildlife if appropriate safeguards are not instituted. This study highlights the need for evidence-based planning for large carnivore conservation across beyond protected area boundaries.

## Acknowledgements

We would like to thank West Bengal Forest Department for providing the permit to conduct the exercise within the protected areas. We would like to thank Anshu Yadav (IFS), Mridul Kumar (IFS), and all the staff of Gorumara Wildlife Division and Jalpaiguri Territorial Division for providing the necessary logistic support during the fieldwork. We would like to thank the authorities and residents of all the tea estates where we conducted the study. We would like to thank Centre for Wildlife Studies and Wildlife Conservation Society-India for providing institutional support. Support for this study was provided by Rufford Foundation (Grant no: 33632-D to AK) and Department of Science and Technology, Government of India (INSPIRE Fellowship, Grant no: IF160391 to AK and INSPIRE Scholarship for Higher Education, Registration No: 201700022793 to AP). We would like to thank Priyanka Das, Motahar Rahaman, and Ramesh Mahali for helping with the fieldwork. We thank Dr. Koustubh Sharma and Dr. Devcharan Jathanna for their help with the analysis. We would also like to thank Dr. Varun R Goswami for his comments and suggestions on the manuscript.

