## Supplementary Files for "Rosettes in a matrix: Predicting spatial variation in density of a large felid in a forest-production mosaic"

### 1 Supplementary Material

#### 1.1 Supplementary Figures

Color should be used for **Supplementary Figure S1, S4 and S5** in print.

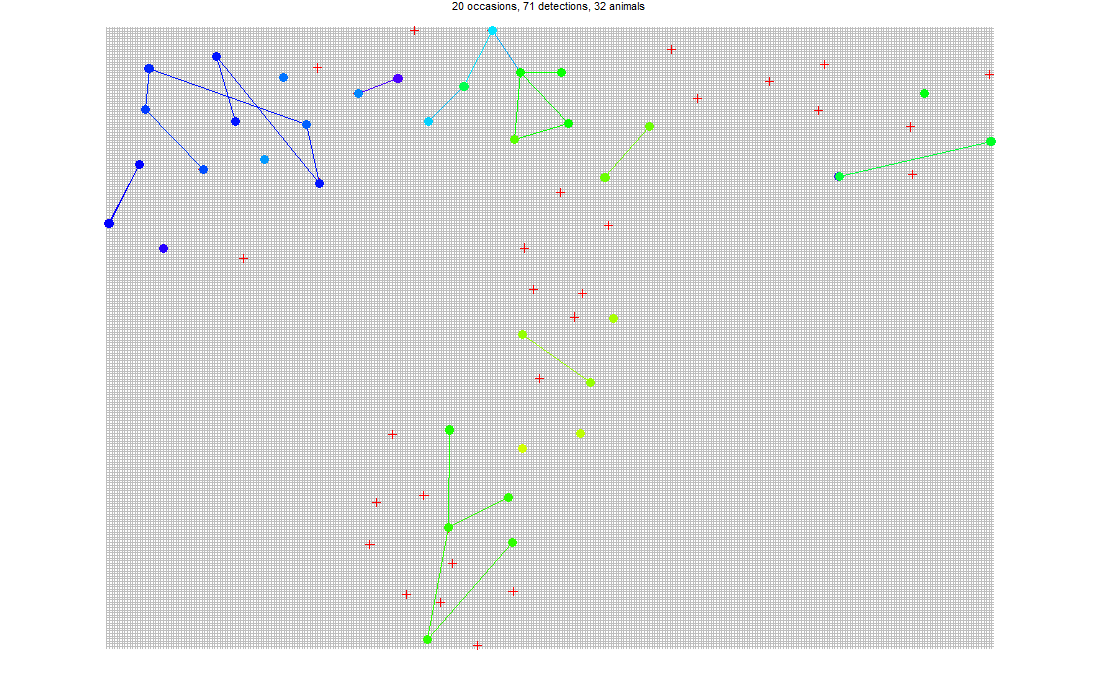
**Supplementary Figure S1.** Arrangement of camera traps, locations of capture and recapture, and the tracks of leopards detected in camera traps. Red crosses represent each camera trap is represented by red crosses; coloured dots represent unique leopard individuals, and the lines represent the tracks of leopard movement from one trap to the other.

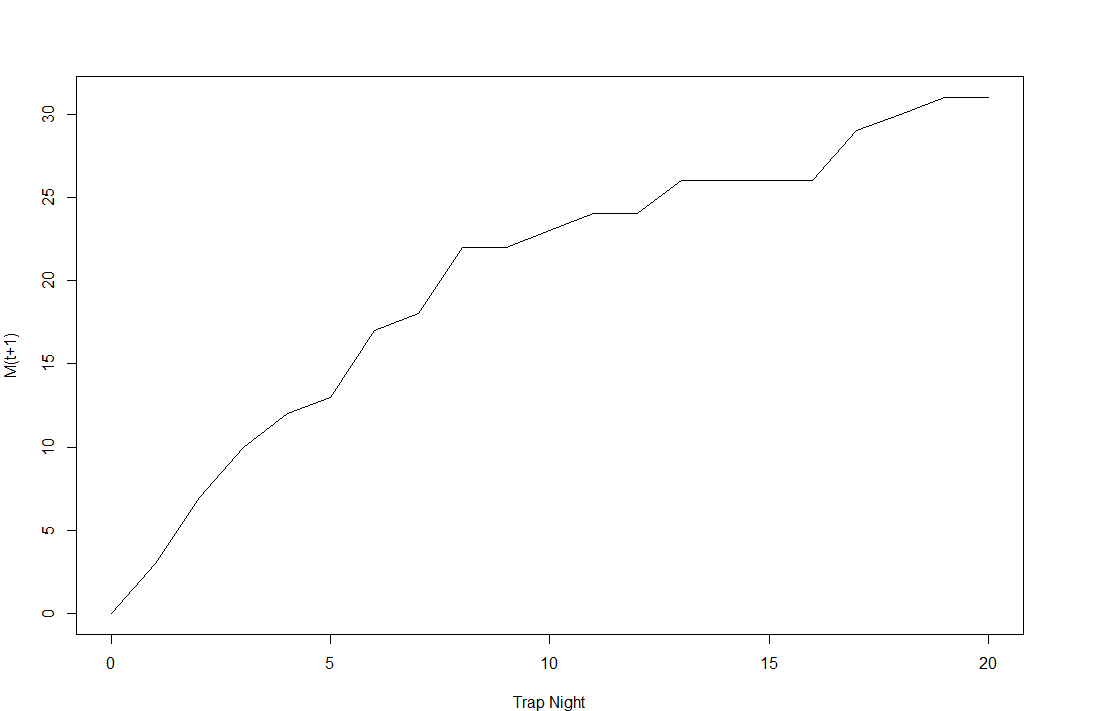

**Supplementary Figure S2.** Plot Showing Cumulative Number of Captured Individuals in each Trap Occasion. The plot does not reach an asymptote suggesting that not all the leopards in the landscape have been photo captured during the sampling period.

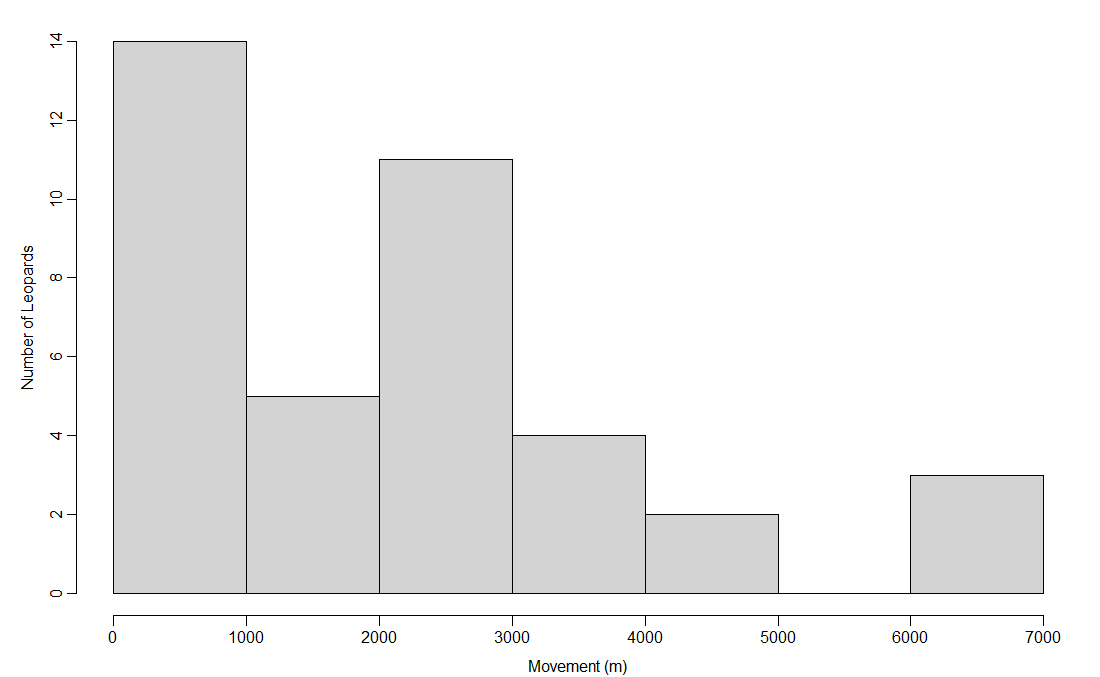

**Supplementary Figure S3.** Histogram showing trap revealed distance of leopard movement across the sampling area. 77% of all the leopard movement occurred within a range of 0-3000 meters from the camera traps. Only 7.7% of all the movement occurred beyond the range of 6000 meters.

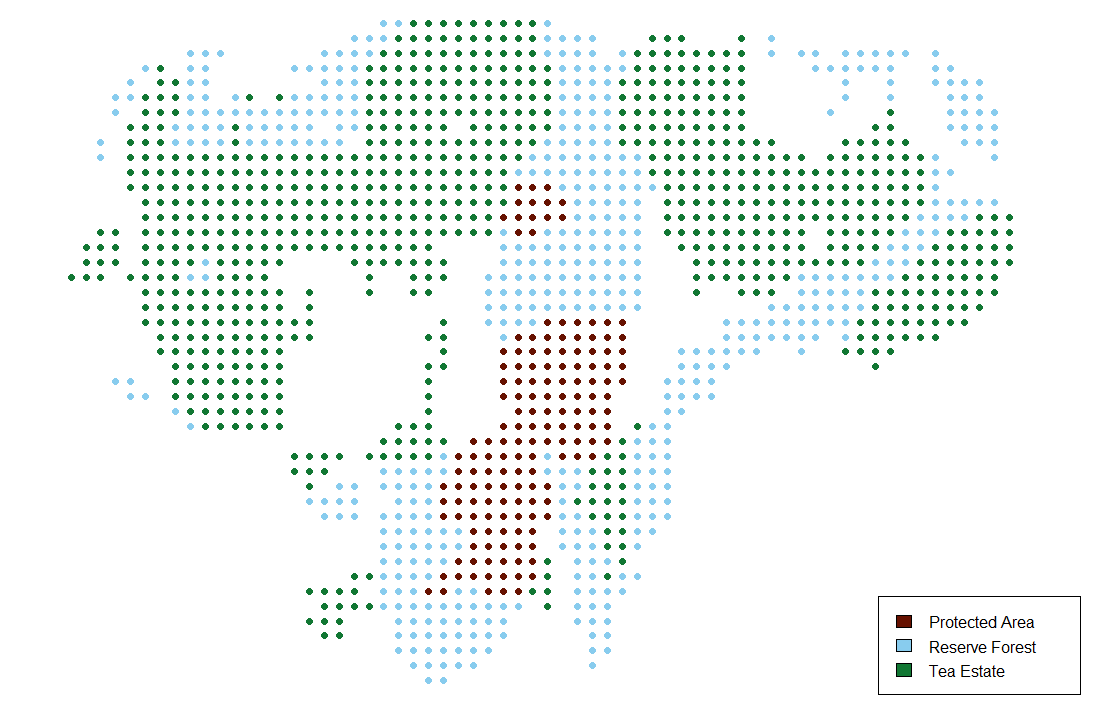

**Supplementary Figure S4.** 6075 meters buffer around camera trap locations forming a state space over which leopard density was estimated. The non-habitat areas were removed to obtain the final habitat mask layer. Habitat Types were added as a categorical covariate. The Reserve Forest and Protected Area together represent the forest habitat in the landscape

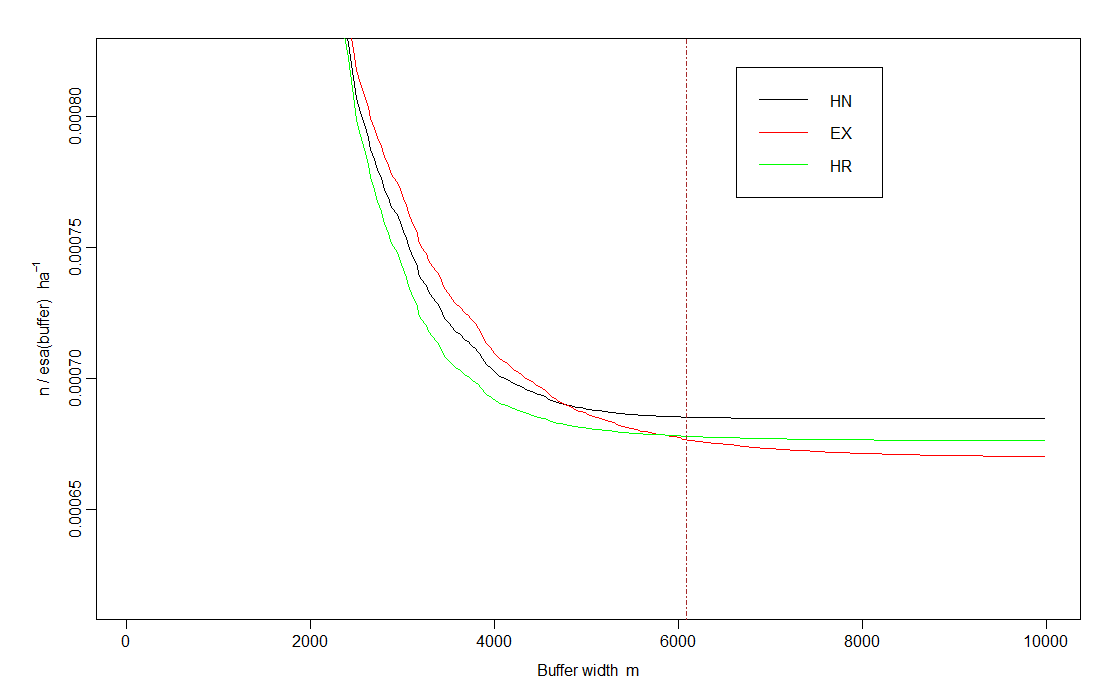

**Supplementary Figure S5.** Stabilization of density with respect to increasing buffer sizes. Half normal detection function hits a plateau comparatively earlier than Hazard rate (HR) and Exponential (EX) detection function. The brown line represents the buffer size used for density estimation

#### 1.2 Supplementary Tables

**Supplementary Table S1.** Capture history file of leopard camera trapping exercise

Object class	capthist

Detector type	count

Detector number	63

Average spacing	1840.075 m

x-range	666484.7 700765.1 m

y-range	2956330 2980032 m

Usage range by occasion

|  | 1 | 2 | 3 | 4 | 5 | 6 | 7 | 8 | 9 | 10 | 11 | 12 | 13 | 14 | 15 | 16 | 17 | 18 | 19 | 20 |
| --- | --- | --- | --- | --- | --- | --- | --- | --- | --- | --- | --- | --- | --- | --- | --- | --- | --- | --- | --- | --- |
| min | 0 | 0 | 0 | 0 | 0 | 0 | 0 | 0 | 0 | 0 | 0 | 0 | 0 | 0 | 0 | 0 | 0 | 0 | 0 | 0 |
| max | 1 | 1 | 1 | 1 | 1 | 1 | 1 | 1 | 1 | 1 | 1 | 1 | 1 | 1 | 1 | 1 | 1 | 1 | 1 | 1 |

Counts by occasion

|  | 1 | 2 | 3 | 4 | 5 | 6 | 7 | 8 | 9 | 10 | 11 | 12 | 13 | 14 | 15 | 16 | 17 | 18 | 19 | 20 | Total |
| --- | --- | --- | --- | --- | --- | --- | --- | --- | --- | --- | --- | --- | --- | --- | --- | --- | --- | --- | --- | --- | --- |
| n | 3 | 5 | 4 | 2 | 2 | 4 | 2 | 6 | 1 | 1 | 3 | 4 | 2 | 3 | 0 | 3 | 3 | 3 | 3 | 2 | 56 |
| u | 3 | 4 | 3 | 2 | 1 | 4 | 1 | 4 | 0 | 1 | 1 | 0 | 2 | 0 | 0 | 0 | 3 | 1 | 1 | 0 | 31 |
| f | 16 | 7 | 6 | 2 | 0 | 0 | 0 | 0 | 0 | 0 | 0 | 0 | 0 | 0 | 0 | 0 | 0 | 0 | 0 | 0 | 31 |
| M(t+1) | 3 | 7 | 10 | 12 | 13 | 17 | 18 | 22 | 22 | 23 | 24 | 24 | 26 | 26 | 26 | 26 | 29 | 30 | 31 | 31 | 31 |
| Losses | 0 | 0 | 0 | 0 | 0 | 0 | 0 | 0 | 0 | 0 | 0 | 0 | 0 | 0 | 0 | 0 | 0 | 0 | 0 | 0 | 0 |
| detections | 4 | 6 | 4 | 2 | 3 | 4 | 2 | 6 | 1 | 1 | 3 | 4 | 2 | 3 | 0 | 3 | 3 | 3 | 3 | 4 | 61 |
| detectors visited | 4 | 6 | 4 | 2 | 2 | 4 | 2 | 6 | 1 | 1 | 2 | 4 | 2 | 3 | 0 | 3 | 3 | 3 | 2 | 4 | 58 |
| detectors used | 61 | 61 | 61 | 61 | 61 | 60 | 60 | 61 | 61 | 60 | 61 | 61 | 59 | 61 | 59 | 59 | 59 | 60 | 59 | 60 | 1205 |

**Supplementary Table S2.** Summary of AIC value for density estimates calculated using halfnormal (HN), negative exponential (EX) and hazard rate (HR) detection functions

|  | **model** | **detectfn** | **npar** | **logLik** | **AIC** | **AICc** | **dAICc** | **AICcwt** |
| --- | --- | --- | --- | --- | --- | --- | --- | --- |
| HN | D~1 g0~1 sigma~1 | halfnormal | 3 | -161.1481 | 328.296 | 329.153 | 0.000 | 0.5694 |
| HR | D~1 g0~1 sigma~1 z~1 | hazard rate | 4 | -160.7308 | 329.462 | 330.943 | 1.790 | 0.2327 |
| EX | D~1 g0~1 sigma~1 | Exponential | 3 | -162.2046 | 330.409 | 331.266 | 2.113 | 0.1980 |

**Supplementary Table S3.** Density estimates (No./100 km^2^) of leopard using halfnormal (HN), negative exponential (EX) and hazard rate (HR) detection function

| **Det Fn** | **Parameters** | **link** | **estimate** | **SE.estimate** | **lcl** | **ucl** |
| --- | --- | --- | --- | --- | --- | --- |
| HN | D | log | **7.96** | **1.56** | 5.43 | 11.67 |
|  | g0 | logit | 0.038 | 0.008 | 0.025 | 0.057 |
|  | sigma | log | 1851.691 | 197.622 | 1503.076 | 2281.161 |
| EX | D | log | **7.93** | **1.58** | 5.38 | 12.69 |
|  | g0 | logit | 0.091 | 0.024 | 0.055 | 0.149 |
|  | sigma | log | 1196.284 | 164.200 | 915.259 | 1563.596 |
| HR | D | log | **7.90** | **1.55** | 5.40 | 11.57 |
|  | g0 | logit | 0.023 | 0.005 | 0.015 | 0.037 |
|  | sigma | log | 2956.506 | 418.834 | 2242.806 | 3897.317 |

**Supplementary Table S4.** Density Estimates (No./100 km^2^) of Different Habitat Types

| **Habitat** | **Effective Sampling Area (Km^2^)** | **Density** | **SE** | **lcl** | **ucl** |
| --- | --- | --- | --- | --- | --- |
| Protected Area | **82.4** | **9.19** | **4.55** | 3.67 | 23.02 |
| Tea Estate | **416.23** | **11.53** | **2.72** | 7.31 | 18.21 |
| Forested Habitat | **343.4** | **4.67** | **2.07** | 2.03 | 10.72 |
